# Emergence of Artemisinin-based Combination Therapy Resistance Markers in *Plasmodium falciparum* from the Brazilian Tri-Border Region of the Guiana Shield

**DOI:** 10.1101/2025.10.13.682088

**Authors:** Yanka E. A. R. Salazar, Antonio M. Rezende, Maria C. S. B. Puça, Jaime Louzada, Maria E. P. Mascarenhas, Sonja Lagström, Danielle Fletcher, Joseli Oliveira-Ferreira, José P. Gil, Tais N. de Sousa

**Affiliations:** Instituto René Rachou, Fiocruz Minas, Fundação Oswaldo Cruz, Belo Horizonte, Minas Gerais, Brazil; Karolinska Institutet, Solna, Sweden; Universidade Federal de Roraima, Boa Vista, Roraima, Brazil; Agilent Technologies Sweden AB, Sundbyberg, Sweden; Instituto Oswaldo Cruz, Fundação Oswaldo Cruz, Rio de Janeiro, Brazil; Institute of Hygiene and Tropical Medicine, Nova University of Lisbon, Lisbon, Portugal

**Keywords:** Malaria, *Plasmodium falciparum*, Drug resistance, Antimalarials, Artemisinin-based combination therapies, Molecular marker, Next-generation sequencing

## Abstract

*Plasmodium falciparum* malaria remains a public health concern in the Brazilian Amazon, particularly among mobile and remote populations. Antimalarial drug resistance threatens elimination efforts, particularly in the Guiana Shield, which has historically been linked to the spread of resistant parasites. We analyzed 99 *P. falciparum* isolates collected in Roraima State, northern Brazil (2016–2020), using targeted deep sequencing. Variants were assessed in *pfcrt*, *pfmdr1*, and *pfk13*, and copy number variation in *pfpm2* and *pfpm3*, including detection of the *pfpm2/3* hybrid. High coverage enabled the detection of both dominant and minority alleles. Among the 99 isolates, the *pfmdr1*-NFD haplotype was found in 6% (6/99, all likely originating from Venezuela), *pfcrt*-C350R in 7% (7/99), and *pfpm2* and *pfpm3* amplifications in 5% (5/94) and 6% (6/97), respectively, and the *pfpm2/3* hybrid in 4% (4/99). No validated or candidate *pfk13* mutations associated with artemisinin partial resistance were detected. This study provides the first report of the *pfpm2/3* hybrid and the *pfmdr1*-NFD haplotype in Brazil, and confirms the presence of *pfcrt*-C350R in isolates from the tri-border area. These variants, associated with reduced susceptibility to lumefantrine and piperaquine, suggest that parasites in northern Brazil already carry resistance markers that are prevalent in neighboring countries. These results highlight the need for continued genomic surveillance to detect early warning signals, guide treatment policies, and contain the spread of parasites with reduced susceptibility to ACT partner drugs.

## INTRODUCTION

In Brazil, the incidence of *Plasmodium falciparum* malaria has significantly decreased since the introduction of artemisinin-based combination therapies (ACTs), particularly artemether-lumefantrine (AL) as the first-line treatment (1). However, imported *P. falciparum* cases have risen in recent years, particularly among mobile and vulnerable populations in the Brazilian Amazon, such as Indigenous people and gold miners (2). In the Yanomami territory, one of the largest Indigenous groups in the Amazon, malaria has sharply increased in association with the expansion of illegal gold mining, aggravating health inequities and creating conditions for uncontrolled transmission (3). From 2011 to 2023, 418,185 malaria cases were reported in Indigenous villages (25.4% *P. falciparum* + mixed infections) and 152,884 in gold mining areas (4). These trends represent a setback for Brazil’s elimination efforts, especially given intensified cross-border mobility that facilitates the introduction and spread of drug-resistant parasites (5).

Antimalarial drug resistance, including partial resistance to artemisinin (ART) and resistance to partner drugs, is an emerging concern that demands strengthened surveillance and timely interventions. While AL remains the first-line treatment in Brazil and other South American countries, dihydroartemisinin–piperaquine (DHA– PPQ) was occasionally used in French Guiana and is widely self-administered in illegal mining sites through products such as Artecom® (dihydroartemisinin + piperaquine + trimethoprim) (6, 7). These drugs are often taken without supervision or correct dosing, contributing to selective pressure and the risk of resistance parasite emergence (6). Mutations in the propeller domain of the *pfkelch13* gene are the current primary markers of partial resistance to ART, although additional markers or combinations may also contribute (8). Although these mutations have not been detected in Brazil, they have been reported in neighboring countries within the Guiana Shield, raising concerns about cross-border spread in areas of intense mobility (9, 10).

Reduced susceptibility to ACT partner drugs has been associated with treatment failures, including artesunate–mefloquine and DHA–PPQ in Southeast Asia (11). More recently, higher tolerance to AL has been documented in parts of Africa (12, 13). Despite this, lumefantrine (LUM) remains a key component of first-line therapies. LUM response is influenced by polymorphisms in the *pfmdr1* gene, especially the NFD haplotype (N86, 184F, D1246), which is selected under AL pressure and modulates drug transport activity (14). This effect is amplified by increased *pfmdr1* copy number (*pfmdr1×N*) (13). This phenomenon is partly explained by the function of the *pfmdr1* product, P-glycoprotein homologue (Pgh), which is inserted in the membrane of the digestive vacuole (DV) in an inside-out position. There, it exports substrates, including antimalarial drugs, into the DV, thereby reducing their cytoplasmic concentration and contributing to resistance phenotypes, particularly against non-DV centric drugs (15).

By contrast, piperaquine (PPQ) acts similarly to chloroquine (CQ), targeting the digestive vacuole and inhibiting heme detoxification, though its mechanism of action is not fully understood (16, 17). Resistance to PPQ is primarily driven by *pfcrt* variants, with modulation by increased copy number of the plasmepsin II and III genes (*pfpm2* and *pfpm3*), which encode digestive vacuole proteases. Among these, the South America–specific *pfcrt*-C350R mutation has been particularly implicated (6, 18). Moreover, recent findings in Africa have shown that the I356T mutation in *pfcrt* undergoes positive selection, co-occurring with *pfpm3* amplifications after repeated treatment cycles, which reinforces the contribution of this SNP in reducing parasite susceptibility to PPQ (19).

To address the threat of emerging multidrug resistance, we investigated the genetic profile of *P. falciparum* isolates from the Guiana Shield, a region historically recognized as a hotspot for the emergence of antimalarial drug resistance in the Americas (6, 20). This highly dynamic border area experiences intense malaria transmission, mainly driven by a high-risk, mobile population engaged in gold mining activities. At the same time, malaria transmission in the Amazon is markedly heterogeneous, with intense outbreaks occurring alongside areas of lower endemicity and limited acquired immunity. In such contexts, resistant genotypes face little competition and can rapidly expand and become fixed through clonal sweeps, particularly under sustained drug pressure (21). Our findings provide a baseline for molecular surveillance and inform cross-border strategies for malaria control. A major strength of this study is the use of targeted deep sequencing, a high-resolution approach, to detect low-frequency resistant variants that are often missed by conventional methods, thereby reinforcing its value in complex transmission settings.

## RESULTS

### Characteristics of participants and epidemiological context

A total of 99 participants were included in this study, with a median age of 31 years (IQR: 25–40). Most individuals were male (68.7%), and the majority of infections were likely acquired in Venezuela (76.7%), particularly in mining areas, according to travel histories. Sample collection sites included Pacaraima (n = 36; 36.3%) and Boa Vista (n = 63; 63.6%). Although most isolates originated from Venezuela, additional samples likely represent infections from Brazil, French Guiana, Guyana, and Suriname (Table 1).

**Table 1.**
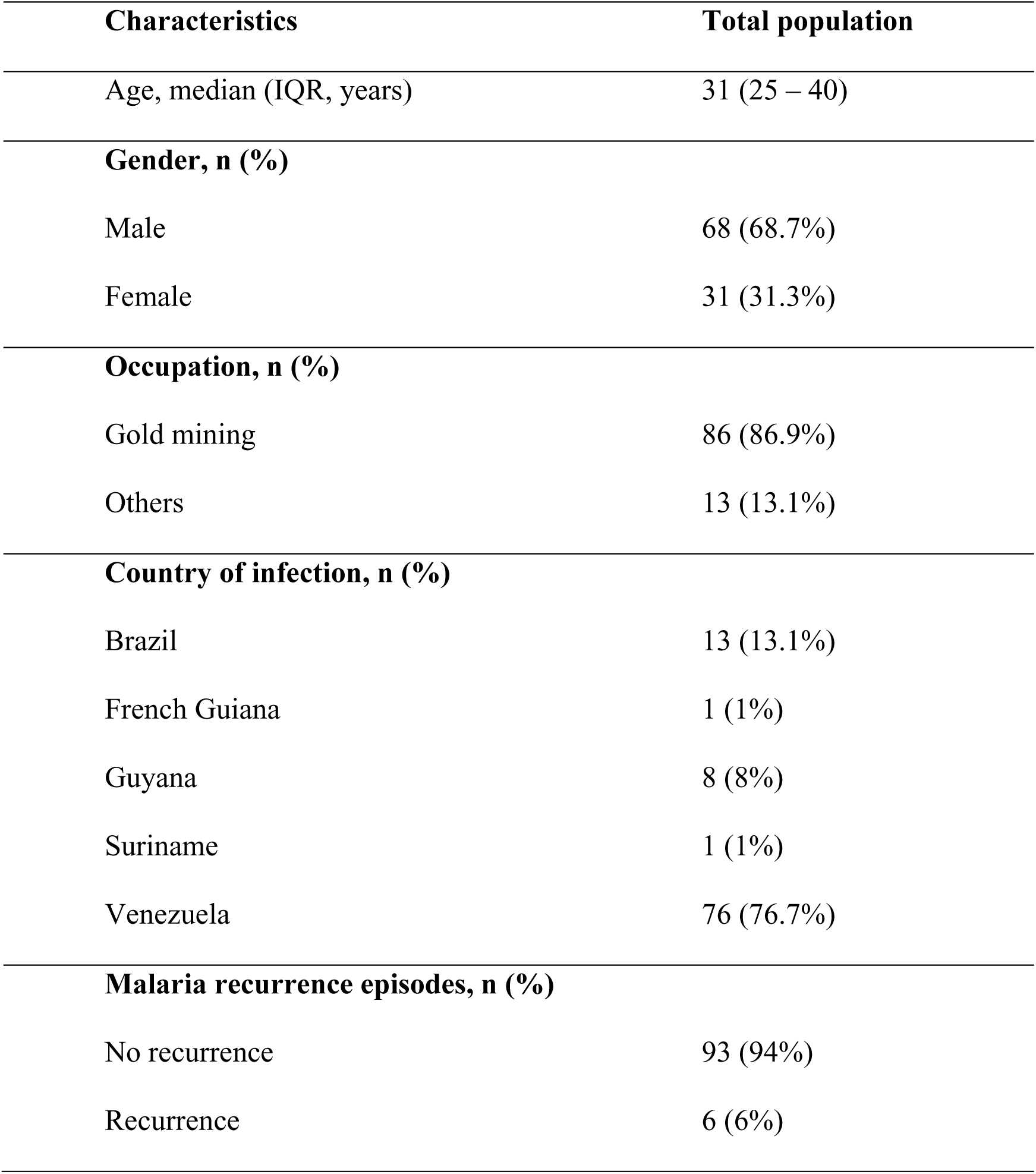
Demographic characteristics of participants included in the study.

### Prevalence of *pfpm2/3* amplification and absence of *pfmdr1* CNV

Copy number variation (CNV) analysis was successful for 97 (98%) samples for *pfmdr1* and *pfpm3*, and 94 (95%) samples for *pfpm2*. No *pfmdr1* amplifications were detected when applying the conservative >1.4 threshold, corresponding to an estimated minimum of 25% of parasites carrying the duplication within an infection (13). Although no isolates reached this cutoff, a few showed values between 1.3 and 1.4, potentially reflecting low-frequency multicopy variants in polyclonal infections. Thus, we cannot rule out the presence of minor *pfmdr1×N* subpopulations below the adopted cutoff.

In contrast, *pfpm2/3* amplifications were detected in 5% of samples (n = 5) for *pfpm2* and 6% (n = 6) for *pfpm3* (Figure 2). Notably, 4% of the samples (n = 4) showed simultaneous amplification of both genes. Additionally, a breakpoint assay targeting the hybrid *pfpm2/3* gene structure detected this variant in the same 4% (n = 4) of isolates, indicating the presence of this structural rearrangement.

**Figure 1.**
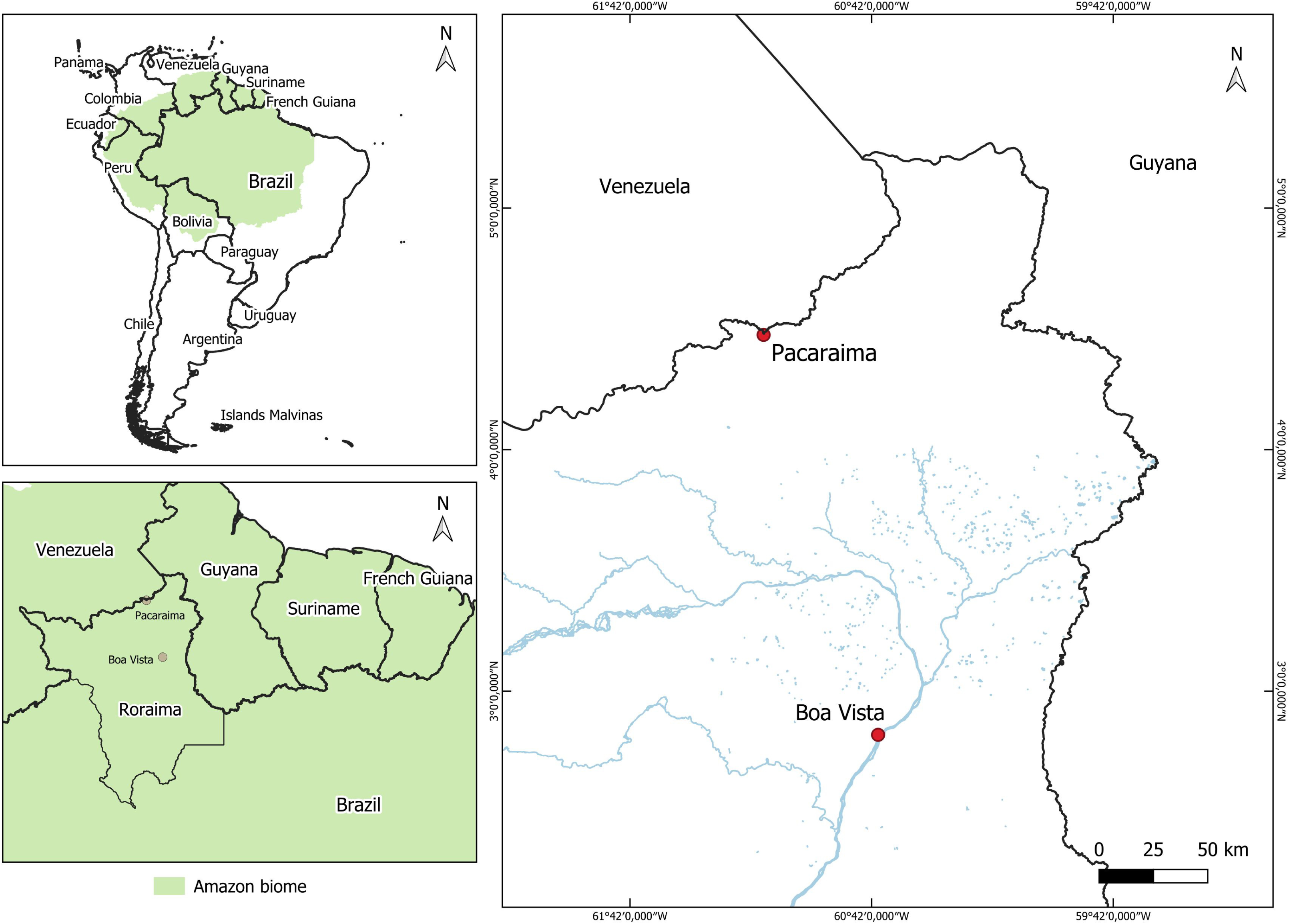
Location of study sites in the Amazon border region of Roraima, Brazil. The map highlights the municipalities of Boa Vista and Pacaraima in Roraima State, northern Brazil. These sites are located near international borders with Venezuela and Guyana, in a region marked by intense cross-border mobility, including migratory flows associated with informal gold mining. The Amazon biome is shaded in green, and red dots identify study sites. This border region plays a critical role in the importation and transmission dynamics of *Plasmodium falciparum* in the Brazilian Amazon.

**Figure 2.**
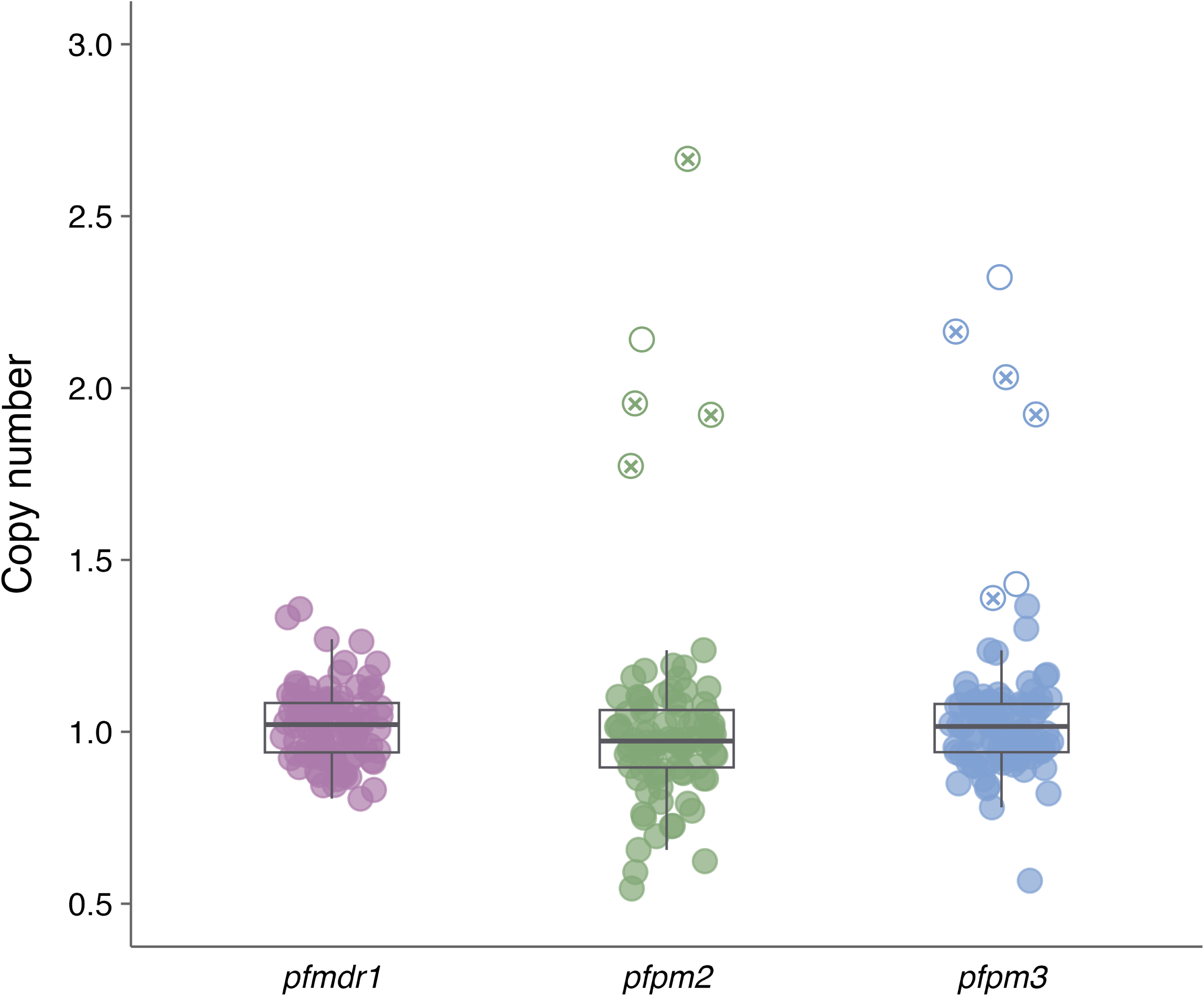
Copy number variation of *pfmdr1*, *pfpm2* and *pfpm3* in *Plasmodium falciparum* isolates. Only samples with relative quantification above 1·4 were considered amplified. Each point represents an individual isolate, with color indicating the corresponding gene. Filled circles represent isolates without copy number amplification, while empty circles indicate amplified copies, and circles marked with x represent a hybrid sequence of *pfpm2/3.* Boxplots show the median (line), interquartile range (box), and values within 1.5× the interquartile range (whiskers).

### NGS Performance and data reliability for key resistance genes

Given the complexity of polyclonal infections, we employed a high-sensitivity targeted sequencing approach to detect minor parasite subpopulations and assessed the quality of NGS data across resistance-associated genes to ensure reliable variant analysis. Sequencing performance was robust, with a mean coverage of 94.7% for the panel of target genes. We herein focused on key genes of clinical and epidemiological importance, *pfcrt*, *pfk13*, and *pfmdr1* genes, which consistently showed high and uniform coverage, supporting the reliability of variant detection. A minimum depth threshold of 100× was established to ensure confident identification of low-frequency variants. The overall sequencing largely exceeded this threshold, averaging more than 1000×. Notably, *pfk13* reached a mean depth of over 2000×, *pfmdr1* 945×, and *pfcrt* approximately 438×. A moderate positive correlation was observed between gene GC content and sequencing depth (r = 0.503; *p* < 0.001; 95% CI: 0.259–0.687), indicating that GC-rich regions tend to yield higher read counts. A weaker but still significant correlation was found between GC content and gene coverage (r = 0.375; *p* = 0.008; 95% CI: 0.104–0.593), suggesting a more limited influence on the breadth of coverage. These results confirm the suitability of the dataset for high-confidence molecular surveillance of antimalarial resistance in *P. falciparum* isolates.

### C350R variant and predominance of *pfcrt* SVMNT 72-76 haplotype

The *pfcrt*-C350R variant was detected in 7% (n = 7) of isolates, with a mean sequencing depth of 262× at this position. Three of these infections were acquired in Brazil, two in Venezuela, one in Guyana, and one in French Guiana. This mutation, previously associated with PPQ resistance, even in the absence of *pfpm2/3* duplications, was not detected in our initial Sanger sequencing but was identified through targeted deep sequencing, which enabled the detection of minority variants.

Classical *pfcrt* polymorphisms at codons 72–76 confirmed the predominance of the SVMNT haplotype, present in most isolates of our dataset (C72S in 84% [n = 80] and K76T in 100% [n = 95]) as shown in Figure 3. In addition, we identified the 7G8 haplotype in 33 isolates (35%), composed of SVMNT, A220S, N326D, and I356L. This haplotype is widely distributed across South America and confers high resistance to CQ and amodiaquine (AQ), while imposing minimal fitness cost to the parasite (25).

**Figure 3.**
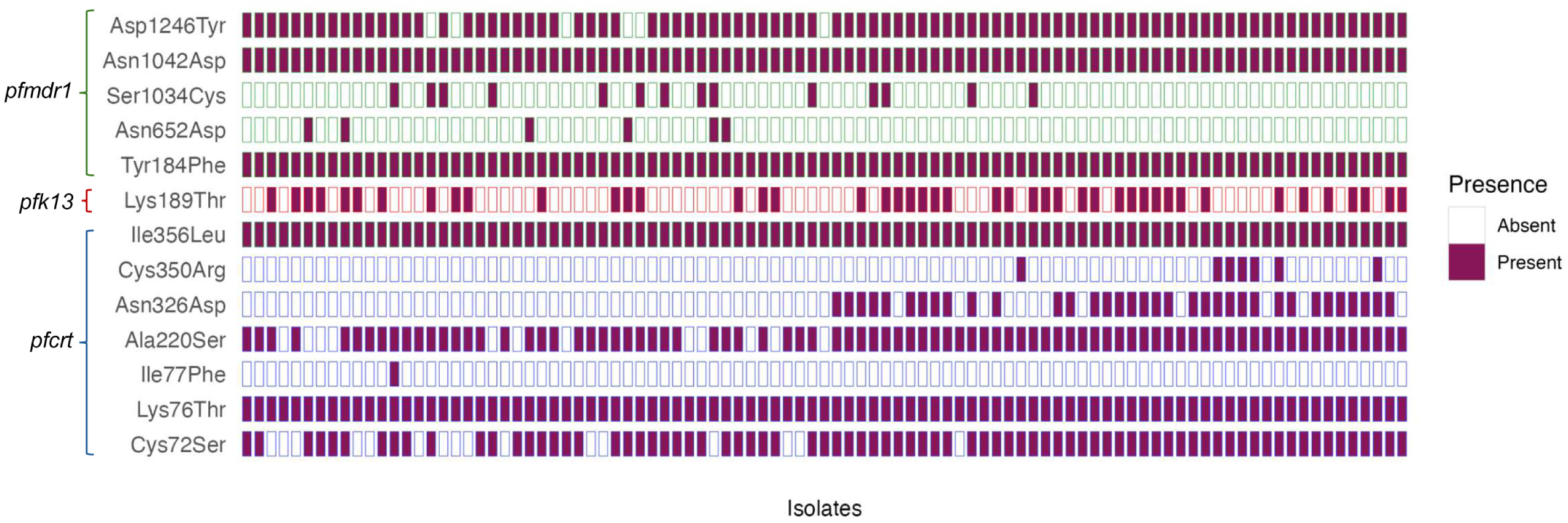
Presence/absence matrix of variants identified in the *pfcrt*, *pfk13*, and *pfmdr1* genes through NGS. The y-axis shows the variants detected in each gene of interest, indicated by color: green (*pfmdr1*), red (*pfk13*), and blue (*pfcrt*). The x-axis represents the analyzed isolates (n = 95). Only variants with a frequency ≥10% of the reads analyzed were considered.

Regarding the *pfk13* gene, no validated or candidate mutations linked to ART partial resistance were detected. The K189T polymorphism was observed in 47% of our dataset (n = 45), although not considered a validated or candidate marker, it has been reported in other studies (26) and is noted here for completeness

### Detection of *pfmdr1* NFD haplotype associated with reduced lumefantrine susceptibility

Analysis of *pfmdr1* revealed that the most prevalent variants were: Y184F (100%, n = 95), N1042D (100%, n = 95) and D1246Y (94%, n = 89), with additional mutations observed at S1034C (15%, n = 14), and N652D (6%, n = 6). The wild-type N86 allele, combined with 184F and D1246, defined the NFD haplotype, which was identified in 6% (n = 6) of isolates (Figure 3). This haplotype has been associated with reduced lumefantrine susceptibility and selection under AL treatment pressure (14). NGS detected five additional isolates carrying 184F that were not identified by Sanger sequencing, consistent with the increased sensitivity of deep sequencing for mixed infections.

### Recurrence analysis

No associations were found between drug resistance markers and recurrence episodes. However, this analysis was limited by the small number of *P. falciparum*-related recurrence cases (n = 6). Recurrence was defined based on cases reported in the SIVEP-Malaria system up to one year before and one year after the date of inclusion in the study. The high proportion of imported infections, mainly from Venezuela, may also have affected recurrence estimates, as some participants likely returned to their country of origin and were not captured by the SIVEP-Malaria system.

## DISCUSSION

The molecular characterization of *P. falciparum* isolates from the Guiana Shield region revealed highly relevant genetic variants with direct implications for antimalarial treatment efficacy in the Amazon. Although many infections were likely imported from neighboring countries, these markers were identified in patients treated in Brazil, underscoring the importance of surveillance in cross-border settings where malaria transmission is shaped by migration, informal drug use, and limited healthcare access. A central concern is the potential introduction and spread of resistant parasites into vulnerable groups, particularly Indigenous populations in remote areas. The Brazil– Venezuela–Guyana tri-border is a hotspot of mobility, driven by gold mining and migration, and overlaps with Indigenous territories. In Roraima, the Yanomami territory has been disproportionately affected, with a sharp increase in malaria linked to illegal mining and poor access to health services (3). In such contexts of heterogeneous transmission and limited immunity, resistant parasites may spread silently and hinder elimination strategies (27).

We identified gene amplifications of *pfpm2* (5%) and *pfpm3* (6%), with 4% of the isolates carrying both. These markers are confirmed determinants of PPQ resistance and have been associated with treatment failures (28). We applied a conservative 1.4 threshold to define gene amplification, based on recent studies suggesting that values above 1.3 may already indicate multicopy subpopulations in polyclonal infections (13, 19). This approach enhances our ability to detect emerging parasite variants that may still be clinically relevant. Although PPQ is not officially used in Brazil, its informal use among gold miners in remote Amazonian areas (2, 6), could impose selective pressure and facilitate the spread of parasites with reduced susceptibility.

We also detected the *pfpm2/3* hybrid sequence in 4% of the isolates, marking the first report of this structural variant in Brazil. Previously described in Southeast Asia, this hybrid arises from a conserved genomic breakpoint between the two *plasmepsin* genes, and has been linked to full PPQ resistance, independent of *pfcrt* mutations (29). While *pfpm2* amplification alone has been associated with treatment failure and *in vitro* resistance (29, 30), the hybrid sequence provides an additional, more specific marker for molecular surveillance (23). The detection of this variant in Brazilian isolates raises concerns about cross-continental introduction through migratory flows, especially given its widespread circulation in Southeast Asia and the increasing movement of individuals between malaria-endemic regions (29, 31–33). Prior studies have already documented similar patterns in the spread of antimalarial resistance, underscoring the risk of global dissemination of resistant genotypes (34–36).

The *pfcrt*-C350R variant, a validated PPQ resistance marker, was found in 7% of isolates, a higher prevalence than previously reported in Brazil (6), suggesting a possible ongoing expansion of this variant. Of these, four were likely imported and three were acquired in Brazil, indicating that local circulation cannot be excluded. Importantly, the variant was identified only through targeted deep sequencing, reinforcing the value of high-resolution tools to capture minority variants. Given that C350R has been associated with reduced PPQ susceptibility even in the absence of *plasmepsin* duplications (6), its presence serves as a strong early indicator of shifting drug response profiles. The combination of multiple PPQ-associated markers increases concern that parasites could, over time, accumulate sets of resistance variants.

No validated or candidate *pfk13* mutations associated with partial resistance to ART were detected, consistent with previous studies in Brazil (37). However, the recent detection of C580Y and G718S in neighboring countries (9, 10), highlights the risk of future introduction into Brazil, given high cross-border mobility.

No confirmed *pfmdr1* amplifications were found above the >1.4 threshold, though some isolates showed intermediate values (1.3–1.4) suggestive of low-frequency multicopy subpopulations. In addition, we detected the *pfmdr1-*NFD haplotype in six isolates, all from patients with infections acquired in Venezuela. This haplotype has been increasingly reported in other endemic regions, particularly in parts of West Africa where AL is widely used as first-line therapy (15, 30, 38, 39), but its role as a validated resistance marker remains uncertain. The detection of both CNV signals and the NFD haplotype highlights the importance of continued monitoring of LUM efficacy in the region. Supporting this, experimental evidence suggests that parasites with the NFD haplotype in a background of increased *pfmdr1* copy number may exhibit enhanced transporter activity, potentially leading to increased drug efflux and reduced drug efficacy (15). A comparable scenario has been reported in Angola, where isolates harboring both variants were associated with declining treatment efficacy of AL (13). These observations raise the possibility that South America could follow a similar trajectory, particularly if AL continues to be used extensively without adequate surveillance to track and contain resistant strains.

This study provides the first report in Brazil of the *pfpm2/3* hybrid sequence and the *pfmdr1*-NFD haplotype, variants linked to reduced susceptibility to PPQ and LUM, respectively. These findings expand our understanding of the diversity and geographic distribution of resistance-associated markers in *P. falciparum* across South America, pointing to the advantages of high-resolution sequencing in detecting minority variants. Beyond their molecular significance, the detection of these variants in a highly mobile tri-border setting, overlapping with Indigenous territories heavily impacted by malaria, reinforces the urgency of coordinated regional surveillance. Overall, our results support genomic surveillance as an essential tool to anticipate emerging resistance, inform locally adapted interventions, and safeguard the efficacy of current antimalarial therapies.

## MATERIALS AND METHODS

### Study Site and Sample Collection

This study was conducted in Roraima state, northern Brazil, with sample collection in the municipalities of Boa Vista and Pacaraima, between March 2016 and March 2020. These sites, located near the Venezuela border, are strategic for malaria surveillance due to high cross-border mobility driven by migration and informal mining (Figure 1) (22). A total of 99 *P. falciparum*-positive samples were included from a previously conducted cross-sectional study in the region (22). Samples with higher parasite densities were prioritized to ensure sufficient DNA quality and quantity for downstream molecular analyses. Participants were symptomatic, aged ≥16 years, non-pregnant, and without signs of severe malaria. The initial diagnosis was based on microscopy, with species confirmation by qPCR. Ethical approval was granted by the Human Research Ethics Committee of the Instituto René Rachou (CAAE: 45337921.0.0000.5091), the Federal University of Roraima Ethical Committee (CAAE: 44055315.0.0000.5302) and Swedish Ethical Review Authority (ref. 2017/499-32). Additional methodological details are provided in the **Supplementary Material.**

### Detection of *pfmdr1* and *pfcrt* SNPs

Polymorphisms in *pfcrt* (K76T, N326S, I356T, and C350R) and *pfmdr1* (N86Y, Y184F, D1246Y) were initially screened using PCR-RFLP and Sanger sequencing, following established protocols (6, 13). These conventional assays served as preliminary analyses to validate marker presence and provided reference points for subsequent next-generation sequencing (NGS). The *pfmdr1*-NFD haplotype was defined by the combination of N86, 184F, and D1246. All preliminary laboratory experiments were performed at Karolinska Institutet (Solna, Sweden). Additional methodological details are provided in the **Supplementary Material**.

### Molecular analysis of *pfpm2, pfpm3* and *pfmdr1* genes

Copy number variation (CNV) of *pfpm2* (PF3D7_1408000), *pfpm3* (PF3D7_1408100), and *pfmdr1* (PF3D7_0523000) was assessed by quantitative PCR (qPCR), using β-tubulin as an internal reference gene. A multiplex qPCR approach was used for *pfpm3* and *pfmdr1* (13), while *pfpm2* CNV followed the method by Pernaute-Lau and colleagues (19). Reference controls included 3D7 (single copy), Dd2 (*pfmdr1* duplication), and NHP1034 (*pfpm2/3* duplication). The *pfpm2/3* hybrid was detected using a validated PCR-based breakpoint assay (23). Relative gene copy number was calculated using the ΔΔCt method, and a conservative threshold of >1.4 was applied to define gene amplification, acknowledging that values between 1.3 and 1.4 may represent minor multicopy subpopulations (13).

### Selective Whole Genome Amplification (SWGA)

All samples underwent SWGA to enrich *P. falciparum* DNA before library preparation and NGS, following the MalariaGEN protocol v2.0 and employed ten previously validated primer pairs specific to the *P. falciparum* genome (24). Amplified products were purified using AMPure XP magnetic beads (Beckman Coulter), with 20 μL of input sample per reaction. Post-purification, DNA concentrations were reassessed using a fluorometric method (Qubit, Thermo Fisher Scientific) to verify the quantity, integrity, and overall suitability of the DNA for sequencing.

### Targeted Deep Sequencing Workflow and Bioinformatic Analysis

Genomic libraries were prepared using a custom hybrid-capture panel and sequenced on the Illumina NextSeq® 2000 platform at the National Genomics Infrastructure, SciLifeLab (Stockholm, Sweden). Library construction included pre- and post-capture amplification steps. Sequencing generated ultra-deep coverage (mean >1000× across target genes), enabling reliable detection of both dominant and minor variants.

Raw reads underwent quality control, adapter trimming, and UMI processing using standard tools, including FastP and FastQC. High-quality reads were aligned to the *P. falciparum* 3D7 reference genome using BWA-MEM, and variant calling was performed with Mutect2 (GATK4), followed by rigorous filtering to retain only high-confidence variants. Identified variants were functionally annotated using SnpEff, and coverage metrics for target genes were calculated using BEDTools. While the overall proportion of polyclonal infections was not formally estimated, the ultra-deep coverage and allelic depth distributions allowed the confident identification of minor alleles. Alleles observed at lower frequencies were retained when supported by high read depth, providing strong evidence that they represented true minority variants within infections. A custom Python script was used to generate summary statistics and visual representations of read depth. Detailed protocols and parameter settings are provided in the **Supplementary Material.**

## AKNOWLEDGMENTS

The authors thank all the patients who participated in this study. The authors thank the Karolinska Institutet Gene Core Facility and the National Genomics Infrastructure at the SciLifeLab. M.C.S.B.P. thanks René Rachou Institute for scholarship support. M.C.S.B.P., M.E.P.M. and Y.E.A.R.S. thank the Coordenação de Aperfeiçoamento de Pessoal de Nível Superior - Brasil (CAPES) for scholarship support (Finance Code 001) and the Programa de Pós-Graduação em Ciências da Saúde do Instituto René Rachou. M.C.S.B.P. and Y.E.A.R.S. thank the Programa PrInt-Fiocruz-CAPES for scholarship support. We also thank the Federal University of Roraima (UFRR) for providing laboratory facilities and logistical support, which were essential for the development of this work.

This work was supported by Fundação de Amparo à Pesquisa do Estado de Minas Gerais (APQ-00516-21), Swedish Research Council Grant (Grant ref. 200075/2022-5), Conselho Nacional de Desenvolvimento Científico e Tecnológico (CNPq) and Programa PrInt-Fiocruz-CAPES. T.N.S. and J.O.F. are recipients of CNPq Research Productivity Fellowships. The funders had no role in the study design, data collection, and interpretation, or in the decision to submit the work for publication.

Conceived and designed the experiments: T.N.S. and J.P.G. Funding acquisition: T.N.S. and J.P.G. Resources: J.L., J.O.F., S.L and D.F. Sample processing and performance of experiments: Y.E.A.R.S., M.C.S.B.P. and M.E.P.M. Data entry and bioinformatics analysis: Y.E.A.R.S. and A.M.R. Writing – original draft: Y.E.A.R.S. and T.N.S. Writing – review and editing: Y.E.A.R.S., A.M.R., J.P.G. and T.N.S. All authors read and approved the final manuscript.

The authors declare no conflicts of interest. Two authors (S.L. and D.F.) are employees of Agilent Technologies. Agilent Technologies provided access to the Magnis NGS Prep System used for library preparation in this study. The company had no role in the study design, data collection and analysis, interpretation of results, or manuscript preparation.

## Data sharing statement

All data generated and analysed in this study are openly accessible through the Supplementary material.

